# 1-Deoxysphingolipids Require Very-Long-Chain Ceramide Synthesis to Induce ER Stress and Neurotoxicity

**DOI:** 10.64898/2025.12.06.692689

**Authors:** Colleen Byrnes, Benjamin A. Clarke, Hongling Zhu, Saurav Majumder, Mari Kono, Richard L. Proia

## Abstract

Sphingolipids play key roles in cellular systems both as membrane components and as signaling molecules. Their biosynthesis, which occurs in the endoplasmic reticulum (ER), begins with the condensation of an amino acid, typically serine, and a fatty acyl-CoA. Under certain pathological conditions, alanine can be substituted for serine in the condensation reaction, producing 1-deoxysphingolipids, which lack the 1-hydroxyl group on the sphingoid base. Unlike typical sphingolipids, 1-deoxysphingolipids are unable to accept a head group modification, which alters their metabolic processing and prevents their canonical degradation. The accumulation of these “headless” 1-deoxysphingolipids causes neurotoxicity in various neurological and metabolic disorders. Here, we conducted a genome-wide CRISPR-Cas9 screen to identify pathways leading to 1-deoxysphinganine–induced toxicity in SH-SY5Y cells, a model used to study neurotoxic responses. Our top genetic hits highlighted the pathway involved in synthesizing ceramides with very-long-chain fatty acids (C22–C26). Using CRISPR-Cas9–modified SH-SY5Y cells with loss-of-function (LOF) mutations in the *TECR* or *CERS2* genes—both critical for producing very-long-chain ceramides—we validated that this pathway was essential for 1-deoxysphinganine–mediated toxicity. Furthermore, we demonstrated that the ceramide synthesis pathway is required for 1-deoxysphinganine to trigger ER stress, as evidenced by significantly increased expression of the unfolded protein response in WT, but not *TECR* or *CERS2* LOF mutant, SH-SY5Y cells exposed to 1-deoxysphinganine. Collectively, these findings identify a specific metabolic pathway for 1-deoxysphinganine leading to very-long-chain 1-deoxyceramide production that culminates in ER stress and toxicity. The findings highlight potential therapeutic targets for neuropathological diseases caused by 1-deoxysphingolipid accumulation.

## Introduction

Sphingolipids are a diverse and abundant class of cellular lipids playing key roles both as membrane components and as signaling molecules (1). The *de novo* biosynthesis of sphingolipids begins in the endoplasmic reticulum (ER) with the condensation of an amino acid, typically serine, and a fatty acyl-CoA, usually palmitoyl-CoA, catalyzed by the enzyme complex serine palmitoyltransferase (SPT) (2, 3). This reaction produces the first sphingoid base, 3-keto-dihydrosphingosine, which is then reduced to form dihydrosphingosine (also known as sphinganine). Sphinganine is subsequently modified with fatty acyl-CoAs by a family of six ceramide synthases that reside in the ER. Each ceramide synthase isoform has a distinct preference for fatty acyl chains of specific carbon lengths. The introduction of a double bond into the sphingoid base converts dihydroceramide to ceramide. From ceramide, complex sphingolipids, such as sphingomyelin and glycosphingolipids, are synthesized by the addition of hydrophilic head groups to the 1-hydroxyl moiety of the ceramide. Eventually, these complex sphingolipids are broken down to regenerate the sphingoid base, which can be phosphorylated at the 1-hydroxyl position by sphingosine kinase to form sphingosine 1-phosphate (S1P). S1P is then degraded by S1P lyase, exiting the sphingolipid metabolic pathway.

Under certain pathological conditions, SPT can substitute alanine for serine, leading to the production of 1-deoxysphingolipids, which lack the 1-hydroxyl group on the sphingoid base and are toxic to cells (Fig. 1*A*) (4). In the absence of the 1-hydroxyl moiety, addition of hydrophilic head groups does not occur, and these so-called “headless” sphingolipids cannot undergo subsequent head group–dependent modification to form sphingomyelin or glycosphingolipids. In addition, the absence of the 1-hydroxyl group prevents the sphingoid base from being phosphorylated to form S1P, which is required for the final degradative step of the sphingolipid metabolic pathway.

**Figure 1.**
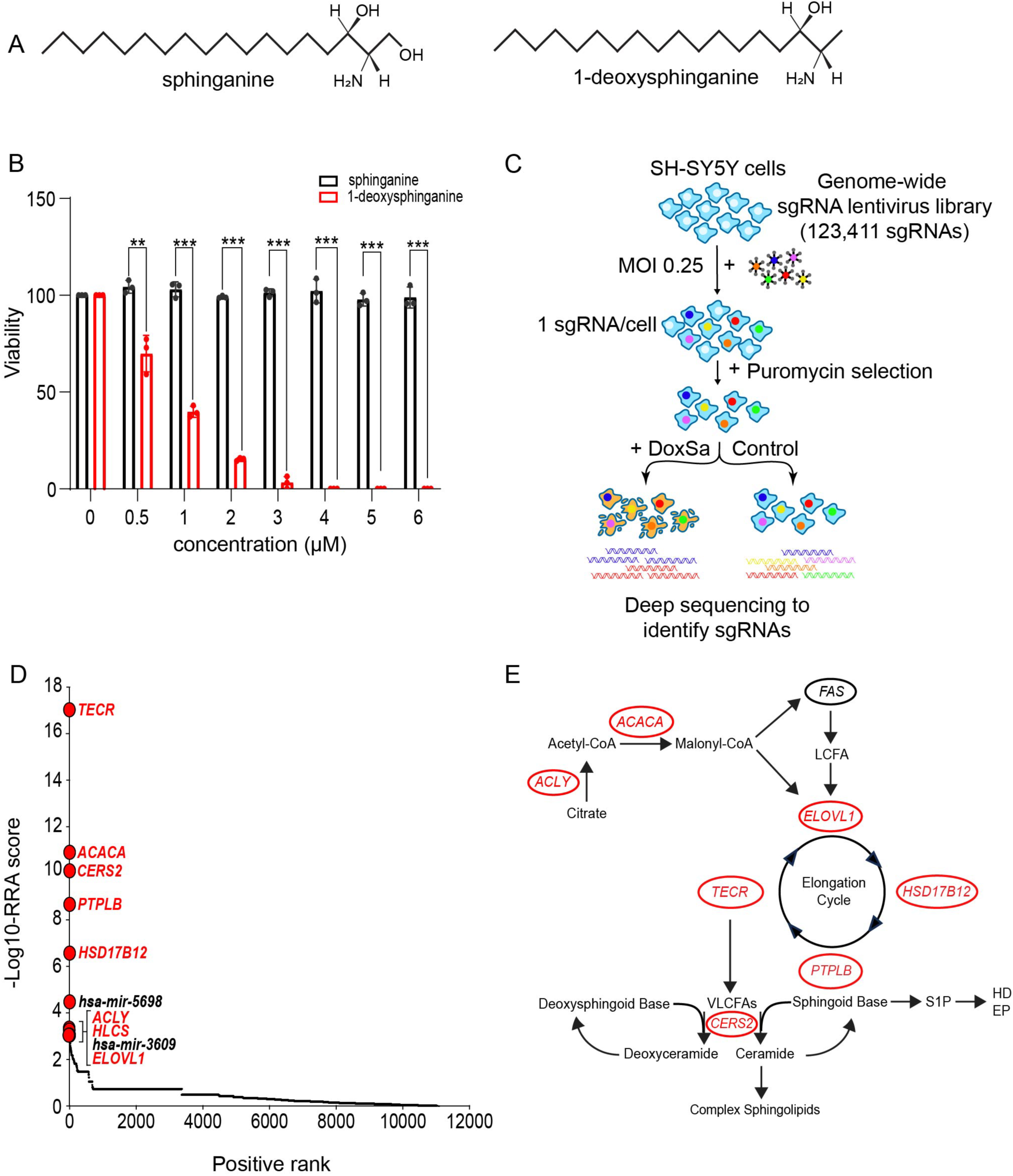
Genome-wide CRISPR/Cas9 screen for genes resistant to 1-deoxysphinganine. (*A*) Structures of sphinganine and 1-deoxysphinganine. (*B*) Viability of SH-SY5Y cells after a 5-day exposure to varying concentrations of sphinganine or 1-deoxysphinganine as determined in a PrestoBlue assay. Data are expressed as mean ± standard deviation (SD) and are representative of three different experiments using the paired *t*-test. **p≤0.01, ***p≤0.001. (*C*) Schematic representation of the strategy for the genome-wide CRISPR-Cas9 screening to identify genes that are resistant to 1-deoxysphinganine in SH-SY5Y cells. MOI, multiplicity of infection; DoxSA, 1-deoxysphinganine; sgRNA, single guide RNA. (*D*) Scatterplot from MAGeCK analysis showing the positively selected gene ranking. The *x* axis represents the positive rank of individual genes and the *y* axis indicates −log_10_ values of corresponding RRA score. The genes involved in lipid metabolism are indicated in red. (*E*) Schematic diagram of very-long-chain ceramide synthesis pathway showing the locations of the positively selected genes. Genes identified in the CRISPR-Cas9 genomic screen are in red. LCFA, long-chain fatty acid; VLCFAs, very-long-chain fatty acids; S1P, sphingosine-1-phosphate; HD, Hexadecenal; EP, Ethanolamine-P.

Elevated levels of 1-deoxysphingolipids have been observed in patients with Hereditary Sensory and Autonomic Neuropathy Type 1 (HSAN1), an autosomal dominant disorder affecting the peripheral nervous system (4–6). In HSAN1, pathogenic variants in the *SPTLC1* or *SPTLC2* genes alter the substrate specificity of SPT, shifting its preference from serine to alanine (7). This substrate shift leads to the accumulation of 1-deoxysphingolipids, which are implicated in the progressive sensory and motor neuropathy, as well as sensorineural hearing loss, characteristic of this disease.

Metabolic disturbances can also elevate the formation of 1-deoxysphingolipids, contributing to neurotoxicity. For example, reduced serine levels as a cause of increased 1-deoxysphingolipids have been observed in macular telangiectasia type 2, a neurodegenerative retinal disease (8, 9). Similarly, serine reductions associated with metabolic disorders may elevate 1-deoxysphingolipid levels, potentially contributing to the peripheral neuropathy commonly seen in type 2 diabetes (9, 10). Furthermore, increased levels of 1-deoxysphingolipids, concomitant with elevated SPT expression, have been linked to peripheral neuropathy induced by the chemotherapy agents paclitaxel and docetaxel (11, 12). These examples illustrate the broad impact of these toxic lipids in neurodegenerative and neurotoxic conditions (5, 13).

To identify the genes responsible for and to deduce the pathway leading from 1-deoxysphinganine, an early biosynthetic metabolite, to the sphingolipid product that causes neurotoxicity, we performed a CRISPR/Cas9 genomic screen in SH-SY5Y cells, a model used to study neurotoxicity (14). We identified and validated the synthesis of very-long-chain 1-deoxyceramides as a cause of cell death and a toxic ER stress response in cells exposed to 1-deoxysphinganine. These results illuminate potential therapeutic targets for disorders caused by the overproduction of neurotoxic 1-deoxysphingolipids.

## Results

### Genome-wide CRISPR/Cas9 screen for genes mediating 1-deoxysphinganine toxicity

1-Deoxysphinganine differs from sphinganine, the normal metabolic intermediate, by the absence of a terminal hydroxyl group at the C1 position (Fig. 1*A*) (4). Because increased levels of 1-deoxysphingolipids have been found to be associated with neuropathologies, we first evaluated their neurotoxic potential in SH-SY5Y neuroblastoma cells (14). SH-SY5Y cells were treated with varying concentrations of either 1-deoxysphinganine or sphinganine, up to 6 μM, and cell viability assessed after 5 days. 1-Deoxysphinganine significantly reduced the viability of SH-SY5Y cells compared with sphinganine, demonstrating toxicity at low micromolar concentrations for the 1-deoxy, but not for the canonical, sphingolipid (Fig. 1*B*).

To identify critical genes mediating 1-deoxysphinganine toxicity, we performed a genome-wide CRISPR/Cas9 screen using the lentivirus-based genome-scale CRISPR/Cas9 Knockout (GeCKO) library, which contains single guide RNAs (sgRNAs) targeting 19,050 genes (15). SH-SY5Y cells stably expressing Cas9 were transduced with the library, selected with puromycin, and grown in the presence or absence of 1 to 4 µM 1-deoxysphinganine for 3 weeks. Genomic DNA from treated and untreated cell pools was then subjected to deep sequencing to identify the relative abundance of sgRNAs (Fig. 1*C*).

The relative abundance of sgRNAs in toxin-treated versus untreated cells was compared using the Model-based Analysis of Genome-wide CRISPR/Cas9 Knockout (MAGeCK) algorithm (16). Genes targeted by individual sgRNAs were ranked based on a modified “robust ranking aggregation” (RRA) score from the MAGeCK analysis (Fig. 1*D*; Table S1). Among the top 10 hits, eight were protein-coding genes involved in lipid metabolism, while the remaining two hits were microRNAs (17, 18). The eight protein-coding genes included ATP citrate lyase (*ACLY*), the primary enzyme responsible for the synthesis of cytosolic acetyl-CoA, a precursor for fatty acid biosynthesis, and acetyl-CoA carboxylase alpha (*ACACA*), a biotin-dependent enzyme that catalyzes the rate-limiting step in long-chain fatty acid biosynthesis. Holocarboxylase synthetase (*HLCS*), the enzyme essential for activating the biotin-dependent enzyme ACACA, was also found (19). Notably, four enzymes involved in the very-long-chain fatty acid elongation cycle—*ELOVL1*, *TECR*, *PTPLB*, and *HSD17B12*—were also identified (17, 18). In addition, ceramide synthase 2 (*CERS2*), which mediates the formation of very-long-chain ceramide species containing fatty acids longer than 22 carbons (20), was among the top 10 hits.

These findings suggest a pathway for 1-deoxysphinganine toxicity that requires the production of very-long-chain fatty acids, which are then incorporated into 1-deoxyceramides by CERS2 (Fig. 1*E*).

### Establishment of cell lines deficient in genes within the very-long-chain ceramide synthesis pathway

To validate the conclusion that the cellular toxicity of 1-deoxysphinganine depends on very-long-chain 1-deoxyceramide biosynthesis, SH-SY5Y cell lines were developed using CRISPR/Cas9–targeted loss-of-function (LOF) mutations in the *TECR* gene (17, 18), which is involved in the fatty acid elongation cycle, or the *CERS2* gene, which is responsible for producing very-long-chain ceramides (20) (Fig. 2*A*). In the *TECR* knockout KO SH-SY5Y cell line, the predominant LOF mutation was a 5-base-pair deletion that introduced a frameshift, resulting in premature stop codon in Exon 6. The predominant mutation in the *CERS2* KO SH-SY5Y cell line was a T insertion that generates a frameshift and a premature stop codon in Exon 2. The CRISPR/Cas9–targeted *TECR* knockout SH-SY5Y cell line exhibited a reduction of approximately 70% in TECR protein levels compared with WT cells, while no CERS2 protein was detected in the CRISPR/Cas9–targeted *CERS2* KO SH-SY5Y cell line (Fig. 2*B*), demonstrating the effect of LOF mutations at each locus.

**Figure 2.**
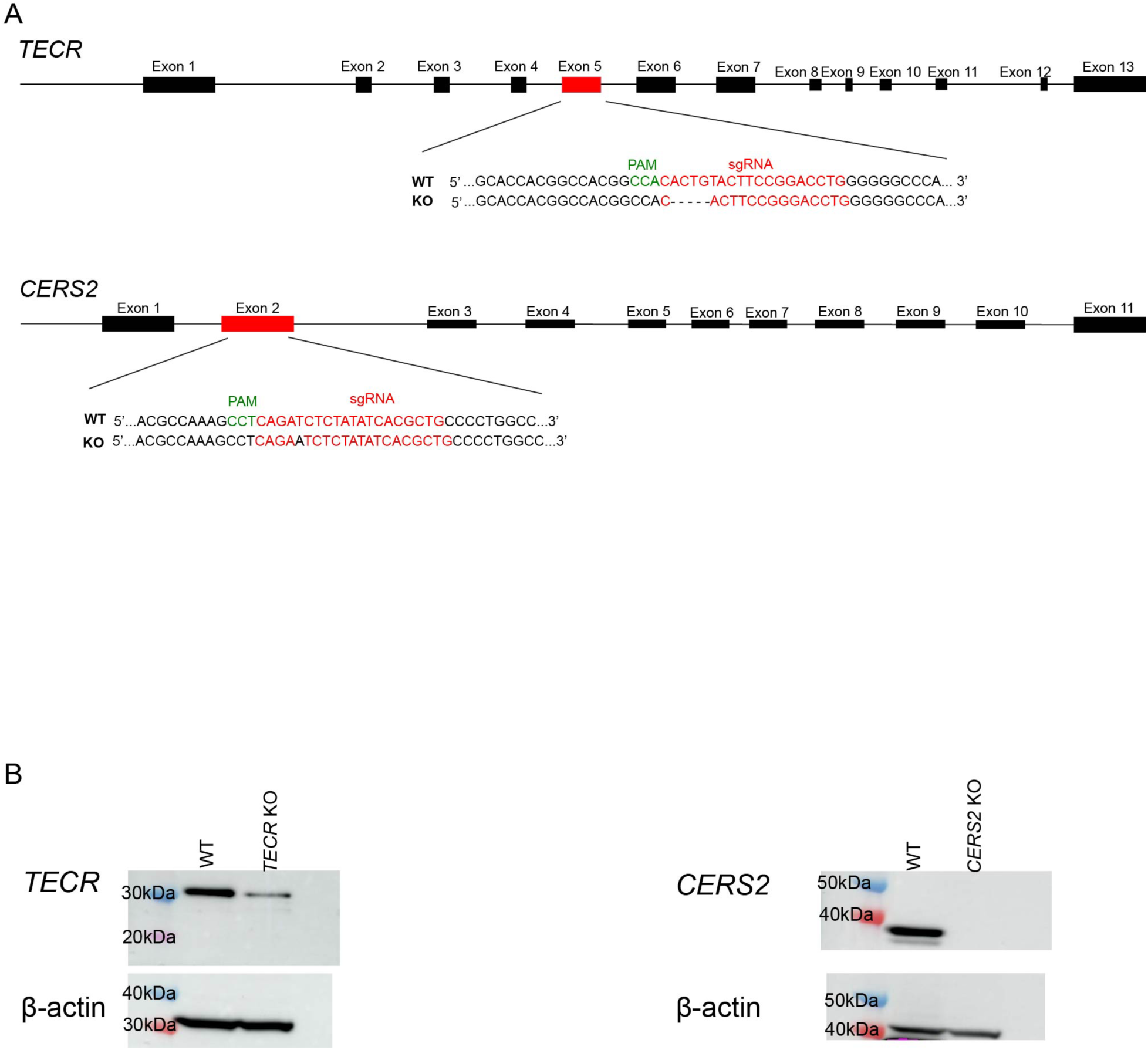
Generation of *TECR* KO and *CERS2* KO SH-SY5Y cell lines. (*A*) Exon/intron structure of the *TECR* and *CERS2* genes, along with sequence and location of the sgRNAs used to create the KO cell lines. Predominant LOF mutations in the *TECR* and *CERS2* genes are shown, (*B*) Western blots of protein expression in the *TECR* (left) and *CERS2* (right) KO SH-SY5Y cell lines. WT SH-SY5Y cells were used as a positive control. β-actin was used as a loading control. Left lane presents molecular weight markers.

### Ceramide levels in TECR and CERS2 KO SH-SY5Y cells

We next used MS analysis to profile the ceramides with varying fatty acid chain lengths produced by WT and mutant SH-SY5Y cells (Fig. 3*A*). In the *TECR* KO cells, there was a significant reduction in the very-long chain-ceramides containing C24 and C24:1 fatty acids, which are the predominant species in WT cells (Fig. 3B). This reduction was accompanied by a concomitant significant increase in shorter chain ceramides, specifically those with C16, C18, and C20 fatty acids. Despite these changes, the total ceramide levels in the *TECR* KO cells were similar to those in WT cells (Fig. 3C).

**Figure 3.**
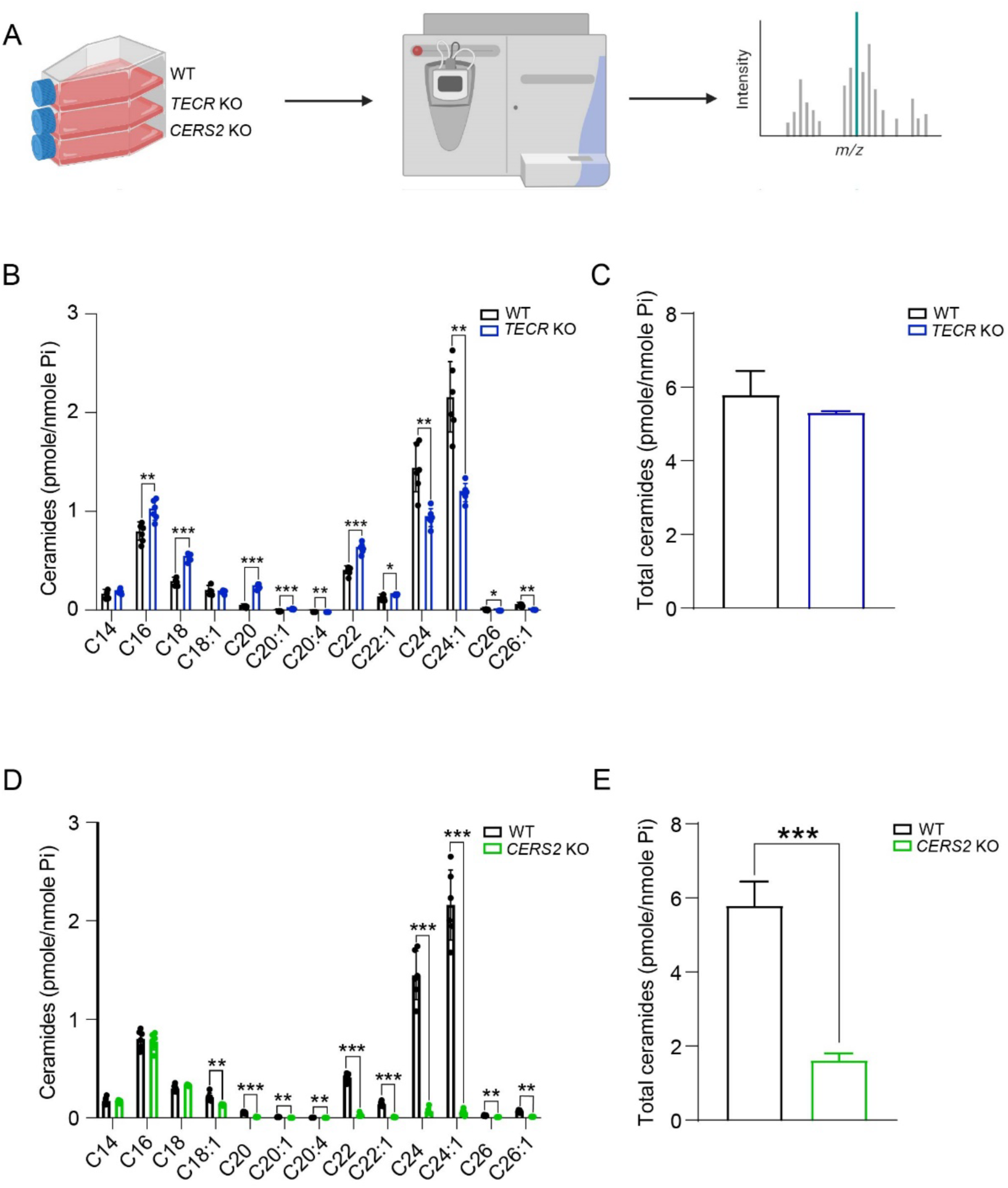
Ceramide levels in *TECR* and *CERS2* KO SH-SY5Y cells. (*A*) Experimental design for ceramide determination by HPLC-MS/MS analysis. Created using BioRender.com. (*B*) Levels of ceramides with different fatty acid chain lengths for *TECR* KO cells compared with WT cells. (*C*) Total ceramides in *TECR* KO cells compared with WT cells. (*D*) Levels of ceramides with different fatty acid chain lengths for *CERS2* KO cells compared with WT cells. (*E*) Total ceramides in *CERS2* KO cells compared with WT cells. Data are expressed as mean ± SD. n=6. Multiple unpaired *t-*test; p-value: * p≤0.05, **p≤0.01, ***p≤0.001.

In contrast, the *CERS2* KO SH-SY5Y cells exhibited a nearly complete depletion of very long-chain-ceramides with C22, C22:1, C24, and C24:1 fatty acids (Fig. 3D). The levels of C14, C16 and C18 fatty acid containing ceramides remained comparable to those in WT cells, while C18:1 and C20 fatty acid containing ceramides were reduced. Total ceramide levels were significantly reduced in the *CERS2* KO cells (Fig. 3E). Thus, loss of *TECR* or *CERS2* compromises very-long-chain ceramide synthesis, but with distinct effects on overall ceramide chain-length composition and total ceramide abundance.

#### 1-Deoxyceramide production in SH-SY5Y cells with an impaired very-long-chain ceramide synthesis pathway

We next assessed the ability of WT and mutant SH-SY5Y cells impaired in the very-long-chain ceramide pathway to convert exogenously added 1-deoxysphinganine into both precursor 1-deoxydihydroceramides and mature 1-deoxyceramides (Fig. 4*A*). In WT cells, the 1-deoxydihydroceramide and 1-deoxyceramide profiles revealed three major very-long-chain forms with fatty acid lengths of C22, C22:1, and C24 (Fig. 4B-E). In the *TECR* KO and C*ERS2* KO lines, all three forms were significantly reduced in the 1-deoxyceramide profiles compared with WT cells (Fig. 4, *C* and *E*). In *CERS2* KO cells, all three forms were also significantly reduced in the precursor 1-deoxydihydroceramide profile (Fig. 4*B*), whereas in *TECR* KO cells, only the 1-deoxydihydroceramide C24:1 form was significantly reduced compared with WT cells (Fig. 4*D*). These results indicate that cells deficient in key genes in the very-long-chain ceramide synthesis pathway also have a reduced capacity to produce both precursor 1-deoxydihydroceramides and mature very-long-chain 1-deoxyceramides.

**Figure 4.**
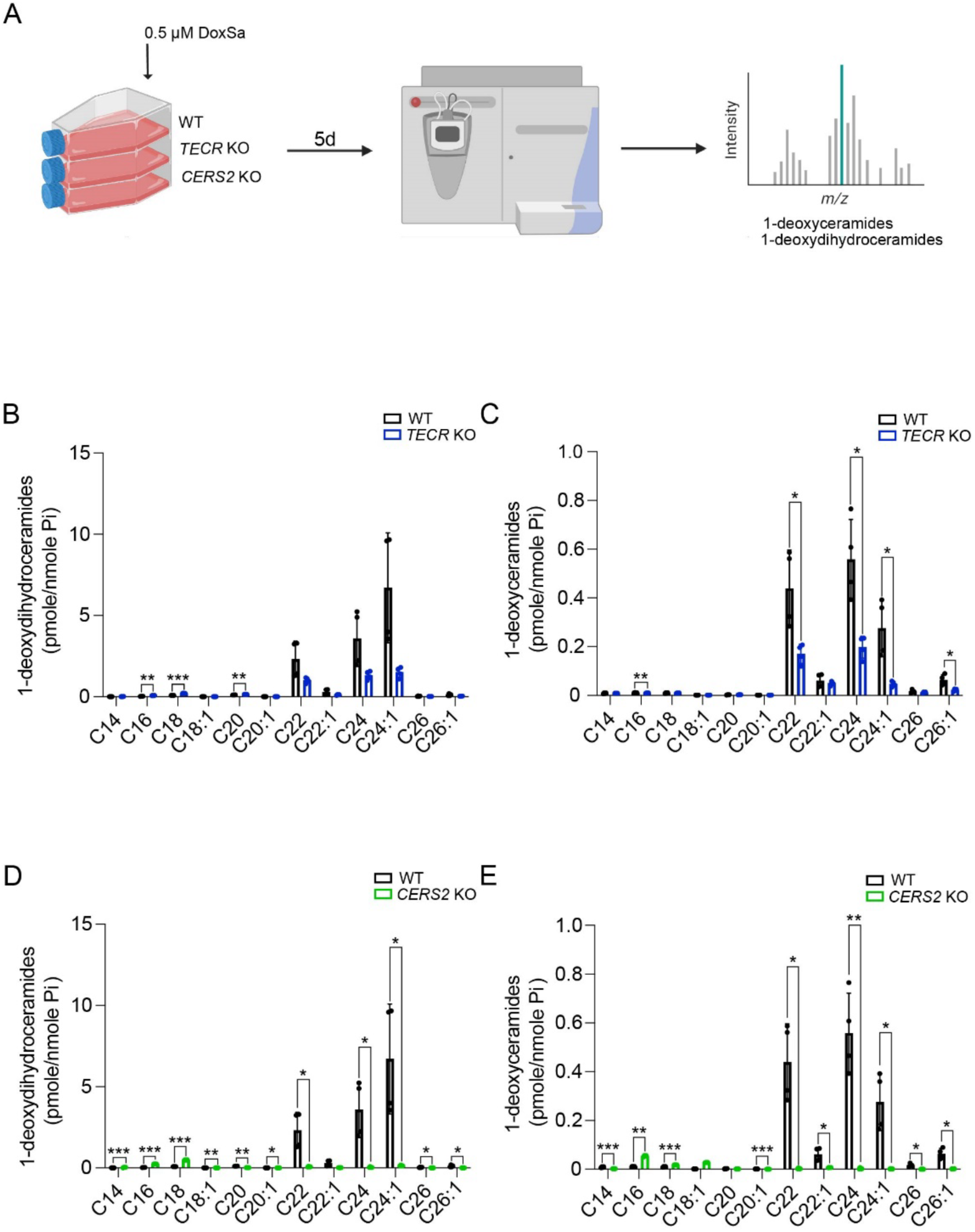
1-Deoxyceramide production in SH-SY5Y cells with an impaired very-long-chain ceramide synthesis pathway. (*A*) Experimental design for 1-deoxydihydroceramide and 1-deoxyceramide determinations by HPLC-MS/MS analysis. DoxSa, 1-deoxysphinganine. Created using BioRender.com. (*B*) Levels of 1-deoxydihydroceramides with different fatty acid chain lengths for *TECR* KO cells compared with WT cells. (*C*) Levels of 1-deoxyceramides with different fatty acid chain lengths for *TECR* KO cells compared with WT cells. (*D*) Levels of 1-deoxydihydroceramides with different fatty acid chain lengths for *CERS2* KO cells compared with WT cells. (*E*) Levels of 1-deoxyceramides with different fatty acid chain lengths for *CERS2* KO cells compared with WT cells. Data are expressed as mean ± SD. n=4. Multiple unpaired *t-*test; p-value: * p≤0.05, **p≤0.01, ***p≤0.001.

### The very-long-chain ceramide synthesis pathway is required for 1-deoxysphinganine toxicity

We next evaluated 1-deoxysphinganine toxicity in *TECR* and *CERS2* KO cell lines compared with the WT SH-SY5Y parental line using a microscopy-based propidium iodide assay to detect cell death (Fig. 5*A*). Cells were treated with 1 μM 1-deoxysphinganine or vehicle (DMSO) for 48 h, then stained with propidium iodide, which labels dead and dying cells, and Hoechst 33342, a nuclear stain. We then quantified the percentage of propidium iodide–positive cells relative to the total number of Hoechst-stained cells. WT cells treated with 1-deoxysphinganine showed significantly increased propidium iodide staining compared with vehicle-treated cells, whereas the *TECR* KO and *CERS2* KO SH-SY5Y cell lines exhibited significantly reduced staining following 1-deoxysphinganine treatment compared with 1-deoxysphinganine–treated WT cells (Fig. 5, *B*–*E*).

**Figure 5.**
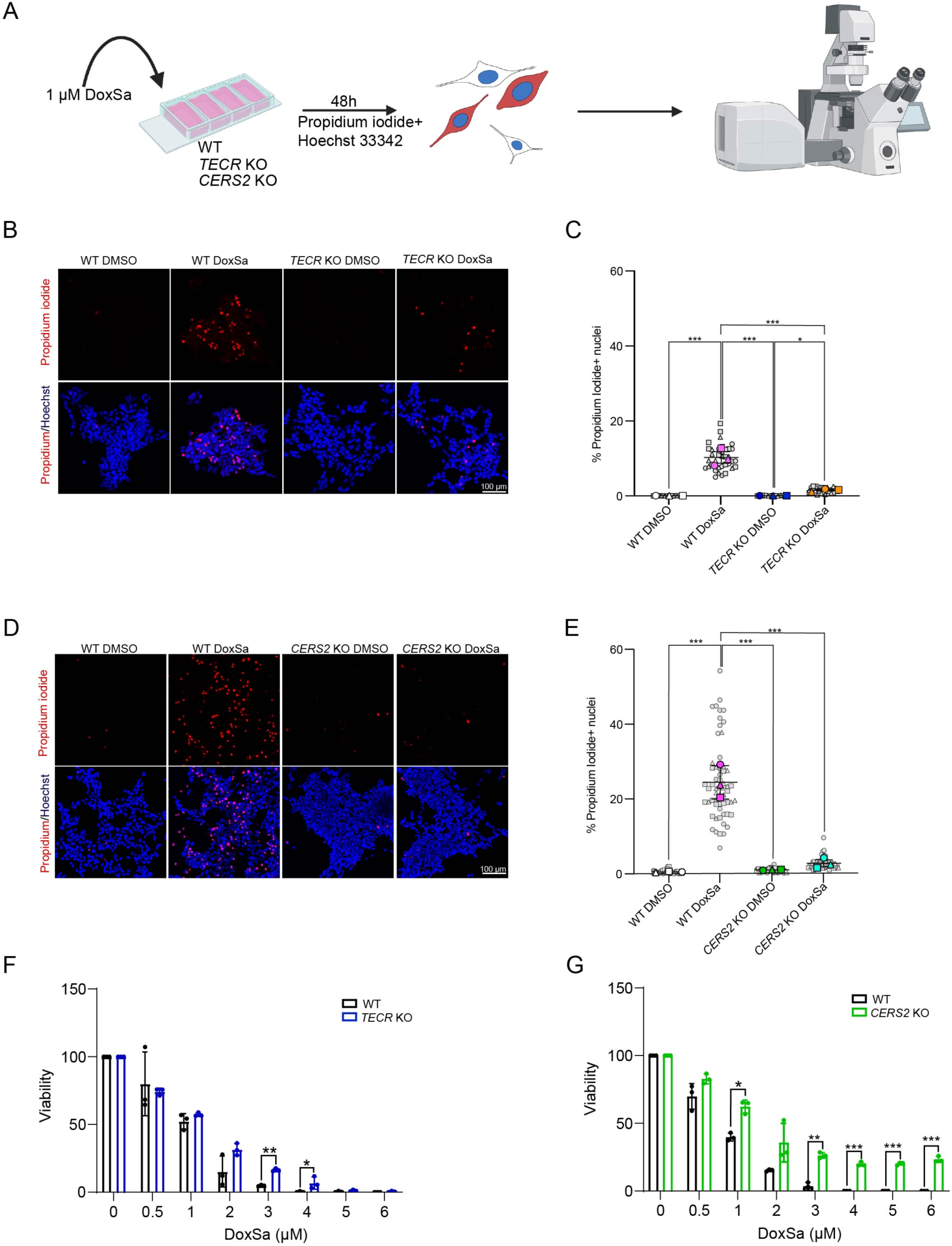
The very-long-chain ceramide synthesis pathway is required for 1-deoxysphinganine toxicity. (*A*) Experimental design of cell death assay. Created using BioRender.com. (*B*) Representative images of WT and *TECR* KO cells treated with or without 1-deoxysphinganine (DoxSa). Propidium iodide staining (dead cells) is in red and Hoechst 33342 staining (nuclei) is in blue. (*C*) Percent of propidium iodide–positive WT cells compared with *TECR* KO cells treated with or without 1-deoxysphinganine. Data are expressed as mean ± SD. One-way ANOVA: *p≤0.05, ***p≤0.001. (*D*) Representative images of WT and *CERS2* KO cells treated with or without 1-deoxysphinganine. Propidium iodide staining (dead cells) is in red and Hoechst 33342 staining (nuclei) is in blue. (*E*) Percent of propidium iodide–positive WT cells compared with *CERS2* KO cells treated with or without 1-deoxysphinganine. Data are expressed as mean ± SD. One-way ANOVA: ***p≤0.001. (*F*) Viability of *TECR* KO cells (left) and *CERS2* KO cells (right) after a 5-day exposure to varying concentrations of 1-deoxysphinganine as determined in a PrestoBlue assay. Gray symbols represent image-level measurements (technical replicates); colored symbols indicate biological replicates. n=3. Data are expressed as mean ± SD and are representative of three different experiments using paired *t*-test. * p≤0.05, ***p≤0.001. Cells without 1-deoxysphinganine were treated with DMSO.

To further confirm the relative resistance of *TECR* KO and *CERS2* KO cell lines to 1-deoxysphinganine, we compared their sensitivity to various 1-deoxysphinganine concentrations with that exhibited by WT cells using the PrestoBlue cell viability assay. Both KO cell lines demonstrated significantly higher resistance to increasing doses of 1-deoxysphinganine than the WT cells (Fig. 5*F*). The increased resistance of the *TECR* KO and *CERS2* KO cell lines to 1-deoxysphinganine supports the conclusion that the very-long-chain ceramide synthesis pathway is critical for the full manifestation of 1-deoxysphinganine toxicity.

### The very-long-chain ceramide synthesis pathway is required for 1-deoxysphinganine–induced ER stress

Previous studies have shown that 1-deoxysphingolipids can induce ER stress and activate unfolded protein response (UPR) genes, including the spliced form of X-box binding protein 1 (XBP1s) and the transcription factor CHOP (8, 21, 22). Chronic UPR activation, particularly through the pro-apoptotic CHOP pathway, can lead to cell death in the setting of sustained, unresolvable ER stress.

To determine whether the very-long-chain ceramide synthesis pathway is required for 1-deoxysphinganine–induced ER stress, we first used a live-cell sensor that reports ER stress by monitoring the IRE1–XBP1 arm of the UPR (23, 24). When the UPR is activated, the XBP1 intron is spliced out, resulting in green fluorescence and an increased green/red fluorescence ratio (Fig. 6A). Treatment of WT SH-SY5Y cells with 1-deoxysphinganine significantly increased this green/red fluorescence ratio, indicating XBP1 intron splicing. In contrast, TECR KO and CERS2 KO cells showed no significant change in the green/red ratio following 1-deoxysphinganine treatment (Fig. 6*B*).

**Figure 6.**
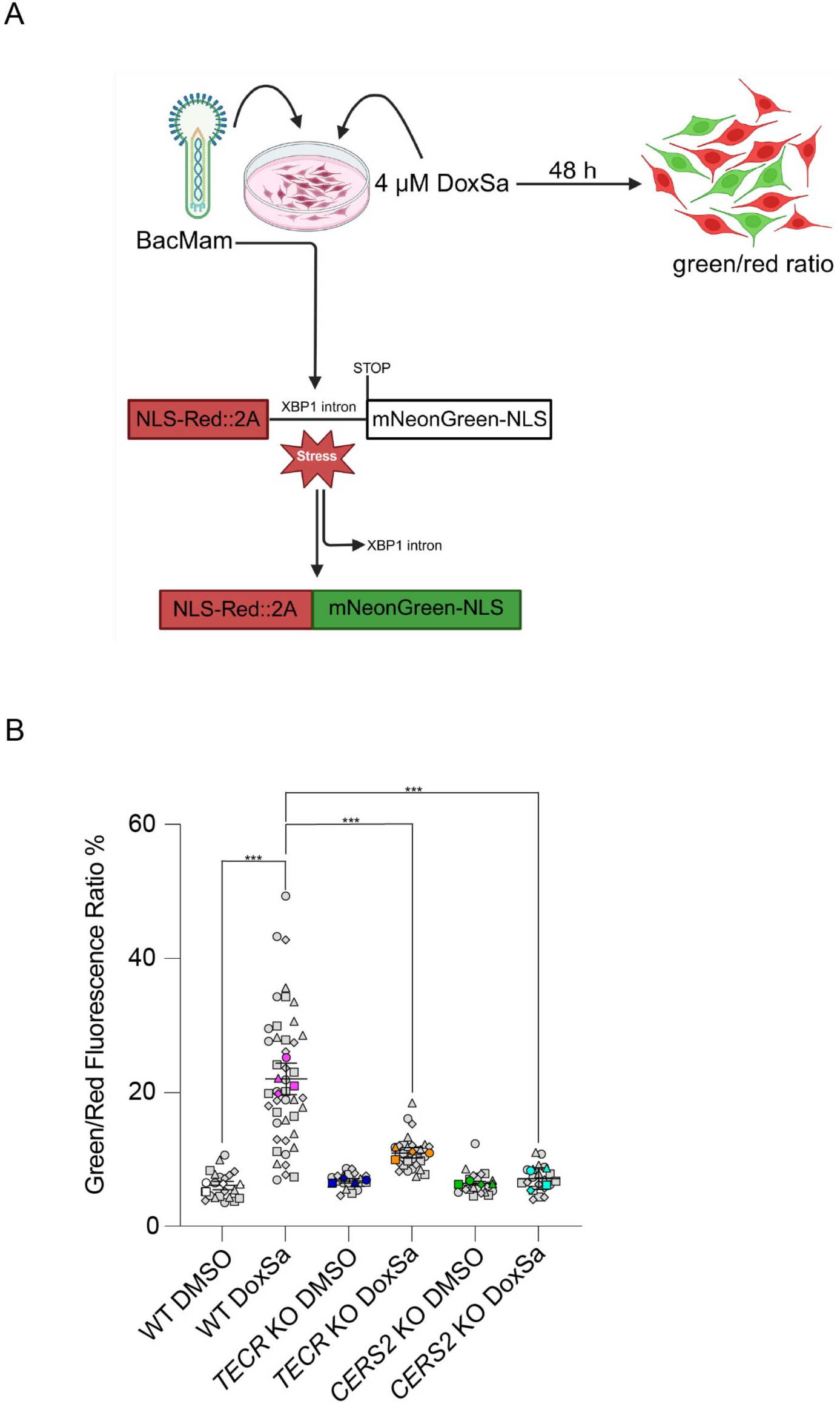
Very-long-chain ceramide synthesis pathway is required for 1-deoxysphinganine–induced XBP1 splicing. (*A*) Experimental design for detecting ER stress using a green/red ratiometric assay. Created using BioRender.com. The BacMam-packaged sensor is based on the splicing of XBP1 mRNA. A red fluorescent protein is constitutively expressed. The sensor produces green fluorescence when the cell undergoes ER stress and triggers the UPR. Cells were treated with or without 4 μM 1-deoxysphinganine (DoxSa) for 48 h and then analyzed for XBP1 splicing. Cells treated without 1-deoxysphinganine were treated with DMSO. (*B*) The green/red fluorescence ratio of WT, *TECR* KO, and *CERS2* KO cells treated with or without 1-deoxysphinganine. Gray symbols represent image-level measurements (technical replicates); colored symbols indicate biological replicates. Data are expressed as mean ± SD. n=3. One-way ANOVA: ***p≤0.001.

To confirm these findings, we performed Western blot analysis for XBP1s, the spliced and active form of XBP1 generated upon UPR activation. XBP1s levels were significantly increased in WT cells after incubation with 1-deoxysphinganine (Fig. 7*A*), whereas *TECR* KO and *CERS2* KO cells did not show a significant increase in XBP1s under the same conditions. These results indicate that WT cells mount an XBP1s-mediated response to 1-deoxysphinganine–induced ER stress, whereas cells lacking the ability to produce very-long-chain ceramides do not.

**Figure 7.**
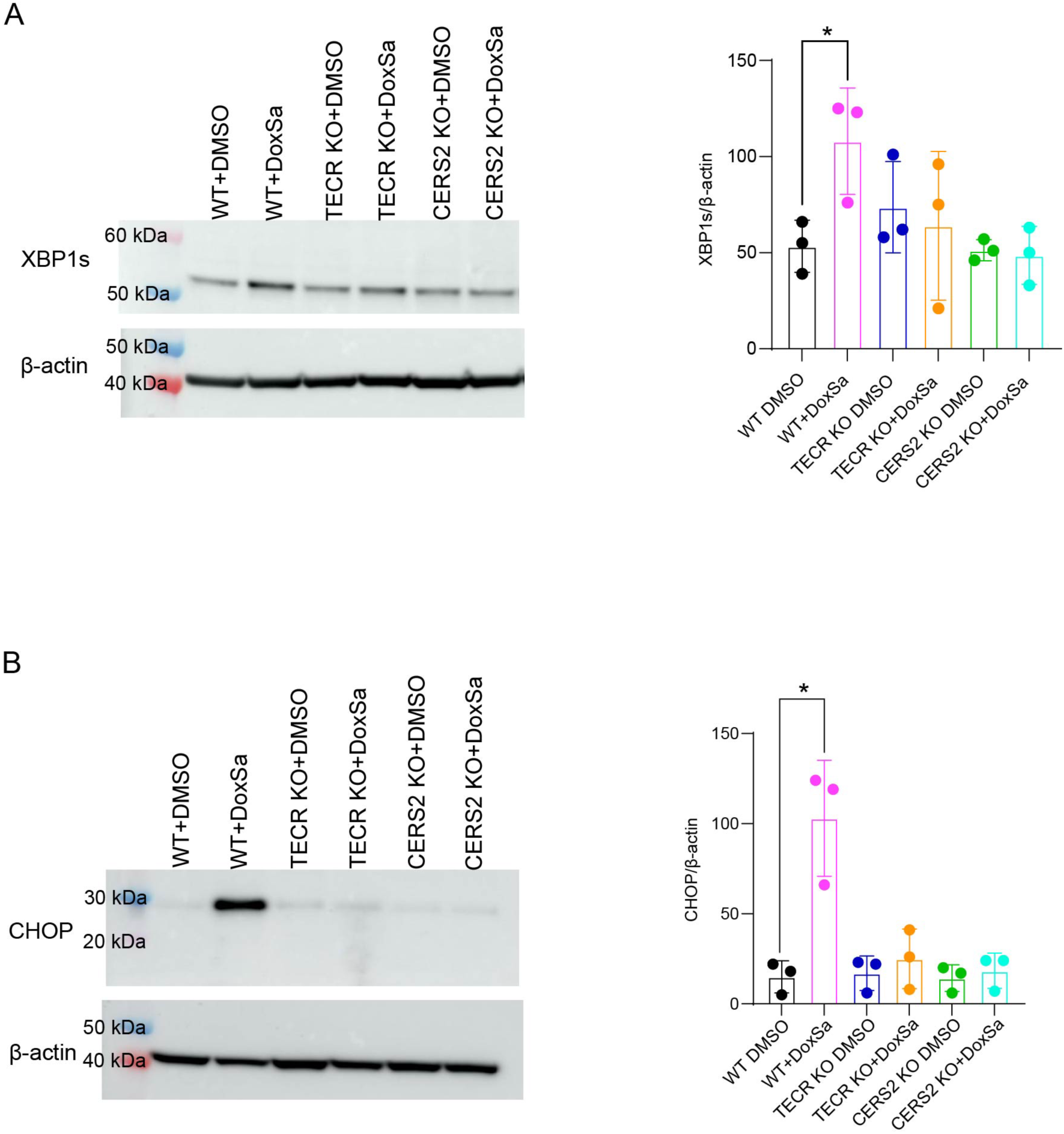
Very-long-chain ceramide synthesis is required for 1-deoxysphinganine–induced XBP1s and CHOP expression. (*A*) Left panel, representative Western blot of CHOP expression in WT, *TECR* KO, and *CERS2* KO cells treated with or without 1 μM 1-deoxysphinganine (DoxSa) for 24 h. β-actin was used as a loading control. Left lane presents molecular weight markers. Right panel, quantification of CHOP protein levels in WT, *TECR* KO and *CERS2* KO with and without 1-deoxysphinganine. Data are expressed as mean ± SD. Each dot represents a separate experiment. n=3. Unpaired *t*-test: *p≤0.05. (*B*) Left panel, representative Western blot of XBP1s expression in WT, *TECR* KO, and *CERS2* KO cells treated with or without 1 μM 1-deoxysphinganine for 24 h. β-actin was used as a loading control. Left lane presents molecular weight markers. Right panel, quantification of XBP1s protein levels in WT, *TECR* KO and *CERS2* KO with and without 1-deoxysphinganine. Data are expressed as mean ± SD. Each dot represents a separate experiment. n=3. Unpaired *t*-test: *p≤0.05. Cells without 1-deoxysphinganine were treated with DMSO.

We next examined CHOP, a pro-apoptotic transcription factor upregulated during prolonged or unresolved ER stress, by Western blotting (Fig. 7*B*). CHOP levels were significantly increased in WT cells following 1-deoxysphinganine treatment but remained unchanged in TECR KO and CERS2 KO cells. The induction of CHOP in WT cells indicates that they experience severe ER stress that can drive a pro-apoptotic program, a response that is absent in the KO cell lines deficient in very-long-chain ceramide synthesis.

Together, these findings demonstrate that the very-long-chain ceramide synthesis pathway is essential for 1-deoxysphinganine to elicit a pro-apoptotic ER stress response.

## Discussion

Excessive production of 1-deoxysphingolipids is thought to underlie neurotoxicity in several conditions, including HSAN1, type 2 diabetes, macular telangiectasia type 2, and chemotherapy-induced peripheral neuropathy (4–6, 8, 9, 11, 12, 25). In this study, we used a genome-wide CRISPR/Cas9 screen to identify genes and pathways contributing to 1-deoxysphinganine–induced toxicity in SH-SY5Y cells. The top hits from the CRISPR-Cas9 screen pinpointed a pathway involved in the production of very-long-chain ceramides, and identified key genes required for the synthesis of very-long-chain fatty acids (17, 18), including *ACLY*, which synthesizes cytosolic acetyl-CoA, a precursor for fatty acid biosynthesis, and *ACACA*, which catalyzes the formation of malonyl-CoA. In addition, the screen detected four enzymes central to fatty acid elongation (*ELOVL1, TECR, PTPLB*, and *HSD17B12*), along with *CERS2*, a ceramide synthase that preferentially incorporates very-long-chain fatty acyl-CoAs (C22–C24) during the formation of ceramide (20). The identification of these metabolic genes suggests that, when it is present, 1-deoxysphinganine is converted into very-long-chain ceramides, ultimately causing toxicity in SH-SY5Y cells. This conclusion was validated in SH-SY5Y cell lines with LOF mutations in *TECR* or *CERS2*, both of which were found to be deficient in very-long-chain ceramide production and showed significant resistance to 1-deoxysphinganine–induced toxicity.

Our conclusion that the very-long-chain ceramide synthesis pathway is essential for 1-deoxysphingolipid cytotoxicity aligns with findings from previous studies. The cytotoxic effects of 1-deoxysphingolipids have been attributed to the accumulation of 1-deoxyceramides, as demonstrated by the inhibition of ceramide synthase with Fumonisin B1 (9, 26–28). Notably, 1-deoxyceramide buildup is linked to anoxic cell death (29), and very-long-chain 1-deoxyceramides (C22–C24) have been associated with neurotoxicity in chemotherapy-induced peripheral neuropathy (11). In addition, studies in yeast indicate that 1-deoxydihydroceramides with long fatty acid chains, such as C26, are significantly more toxic than those with shorter C16 or C18 chains (30), further underscoring the heightened toxicity associated with very-long-chain 1-deoxyceramides.

Earlier studies have also shown that excess 1-deoxysphingolipids can trigger an ER stress response across various cell types (8, 21, 22, 31), contributing to cellular toxicity. In our experiments with *TECR* or *CERS2* LOF SH-SY5Y cells, which lack the ability to produce very-long-chain ceramides, we demonstrated that the production of very-long-chain 1-deoxyceramides specifically induced ER stress (as evidenced by increased expression of XBP1s and the pro-apoptotic transcription factor CHOP) in WT, but not *TECR* or *CERS2* LOF mutant, SH-SY5Y cells exposed to 1-deoxysphinganine. Because these 1-deoxyceramides are synthesized within the ER and cannot be processed through the typical degradation pathway involving S1P lyase, they may accumulate, disrupting ER function and triggering an ER stress response. In yeast, 1-deoxysphingolipids have been shown to cause ER collapse, leading to cell death (30). Similarly, exposure of MEF cells to 1-deoxysphingolipids has been associated with ER abnormalities and cellular toxicity (31). Our findings support the idea that very-long-chain 1-deoxyceramides, in particular, have harmful effects on ER integrity.

Our results suggest that strategies to reduce the activity of the very-long-chain ceramide synthesis pathway may help limit neurotoxicity in diseases marked by excessive 1-deoxysphingolipid production. A potential therapeutic target could be CERS2, which is broadly expressed, including in nervous tissue (20). However, complete inhibition of *CERS2*, as shown in KO mouse models, can lead to toxicity in the liver, kidney, and nervous system (32, 33). Alternatively, therapeutic approaches could focus on modulating the ER stress response, which may alleviate the cellular damage caused by 1-deoxysphingolipid accumulation.

In summary, our findings illustrate the pivotal role of the very-long-chain ceramide synthesis pathway in mediating 1-deoxysphingolipid-induced toxicity in neuronal cells. The accumulation of these 1-deoxyceramides disturbs ER integrity and triggers an ER stress response that culminates in cell death. By targeting these pathways, it may be possible to develop treatments to mitigate the neurotoxic effects of 1-deoxysphingolipid accumulation in various neurological and metabolic disorders.

### Experimental Procedures

#### Genome-wide CRISPR/Cas9 screen

A genome-wide CRISPR/Cas9 KO screen was performed as previously described (15, 34, 35). Human GeCKOv2 CRISPR knockout pooled library was a gift from Feng Zhang (Addgene #1000000048). Briefly, SH-SY5Y cells expressing Cas9 (GeneCopoeia) were transduced with a multiplicity of infection of 0.25 with the human GeCKO v2 library (15) for a total coverage of 50X. Puromycin was added 24 h post-transduction at a concentration of 0.5 μg/ml. After 5 days of puromycin selection, cells were cultured with or without 1-deoxysphinganine (d18:0) (Avanti Polar Lipids, 860493) at a concentration of 1 to 4 μM. After treatment for 3 weeks with 1-deoxysphinganine, the cells were harvested for genomic DNA isolation using the Qiagen Blood & Cell Culture DNA Maxi Kit (Qiagen, 13362) according to the manufacturer’s instructions. Genomic DNA from each group was used as template DNA to amplify the sgRNA region by PCR. The first round of PCR product of the PCRs was used as a template for another round of PCR to amplify and attach Illumina-compatible multiplexing sequence adaptors (34). The libraries were then subjected to single-end sequencing by Illumina NextSeq for control- and 1-deoxysphinganine–treated cells to identify the copy number of sgRNAs. Sequence reads were analyzed using the MAGeCK computational tool (16). The resultant RRA scores are presented in figures by R script.

#### Generation of KO cell lines

Parental SH-SY5Y cells were obtained from ATCC (CRL-2266). SH-SY5Y *TECR* KO and *CERS2* KO pools were generated by Synthego. The cells were cultured in Eagle’s Minimum Essential media (EMEM) (ATCC, 30-2003) containing Ham’s F12 nutrient mix media (Thermo Fisher Scientific, 31765035), 10% fetal bovine serum (FBS), and penicillin-streptomycin (Thermo Fisher Scientific, 15140148). *TECR* KO and *CERS2* KO pools were sorted into a 96-well plate using a BD FACSAria II flow cytometer (BD Biosciences) to generate individual single-cell clones. Genomic DNA was extracted from these cell clones using the DNeasy Blood & Tissue Kit (Qiagen, 69506) following the manufacturer’s instructions. Restriction enzyme sites were identified near the cut sites of the sgRNAs of *TECR* and *CERS2* to allow for screening of indels. The restriction enzyme *BglII* (New England Biolabs, R0144L) was used to determine the absence or presence of indels in *CERS2*. The following primers were used for *CERS2*: forward 5’-CGGGTAGAGGGTGAGAACAAATAA −3’; reverse: 5’-TGTAATACGGTGGCTTTCCAGTTTTAA-3’. *BglII* will not cut if indels are present. The restriction enzyme *TatI* (Thermo Fisher Scientific, ER1291) was used to determine if indels were present in *TECR*. The following primers were used for *TECR*: forward 5’-CTGTAACTGCCCTGTTCTCCC −3’; reverse: 5’-GTCACTCATTCCACTACATCAAGC-3’. *TatI* will not cut if indels are present. The putative KO clones were confirmed by DNA sequencing.

#### Cell culture and cell viability assays

SH-SY5Y cells were cultured in EMEM containing Ham’s F12 nutrient mix media, 10% FBS, and penicillin-streptomycin. A 5 mM stock solution of 1-deoxysphinganine was prepared in DMSO. A 10 mM stock solution of sphinganine (d18:0) (Avanti Polar Lipids, 860498P) was prepared in ethanol. For cell viability assays, WT, *TECR* KO, or *CERS2* KO SH-SY5Y cells were exposed to 0 to 6 μM sphinganine or 1-deoxysphinganine. After 3 days, PrestoBlue Cell Viability Reagent (Thermo Fisher Scientific, A13262) was added to determine cell viability. The cells were incubated in PrestoBlue for 2 h, followed by analysis on the PerkinElmer Enspire plate reader.

#### Western blot analysis

RIPA Lysis and Extraction Buffer (Life Technologies, 89900) was used to extract protein from WT, *TECR* KO, or CERS2 KO SH-SY5Y cells following the manufacturer’s instructions, with the addition of HALT protease inhibitor cocktail (Thermo Fisher Scientific, 78429). Equal amounts of protein (30 μg) were separated on a 4–12% Novex Bis-Tris gel (Thermo Fisher Scientific, NP0321BOX) with the Novex Sharp Pre-stained protein standard (Thermo Fisher Scientific, LC5800) and transferred onto either a nitrocellulose or polyvinylidene fluoride membrane using the iBlot2 Gel Transfer Device (Thermo Fisher Scientific, IB21001). The membranes were blocked in 5% milk for 1 h at room temperature and incubated overnight with primary antibodies at 4°C. The following antibodies were used: CERS2 (Sigma Aldrich, HPA027262); TECR (Thermo Fisher Scientific, A305-515A-T); CHOP (Cell Signaling Technology, 2895S); XBP1s (Cell Signaling Technology, 12782S); and β-actin (Cell Signaling Technology, 5125S). Membranes were washed in 5% milk and incubated with the either horseradish peroxidase–conjugated anti-mouse (MilliporeSigma, AP124) or anti-rabbit (Millipore Sigma, AP132) secondary antibody in 5% milk. Membrane signals were detected using the ECL Prime Western blotting system (MilliporeSigma, GEPRN2232) and imaged on the Amersham Imager 680 (GE Healthcare Life Sciences). β-actin was used as a loading control. The blots were analyzed using the ImageQuant TL 8.2 analysis software (GE Healthcare Life Sciences) and normalized to the β-actin signal.

#### Mass spectrometry

Ceramides, 1-deoxydihydroceramides, and 1-deoxyceramides in cell lines were measured by HPLC-tandem MS (HPLC-MS/MS) by the Lipidomics Core at the Medical University of South Carolina as previously described (36).

#### Cell death assays

WT or *CERS* SH-SY5Y KO cells (4 x 10^4^) were seeded in poly-d-lysine (Millipore, A-003-E) coated μ-Slide 4-well chamber slides (ibidi GmbH, 80426). WT or *TECR* KO SH-SY5Y cells (3 x 10^5^) cells were plated on 35-mm plates (Crystalgen, 191-3000). The next day, the cells were treated with vehicle (DMSO) or 1 μM 1-deoxysphinganine. After 48 h, propidium iodide (Thermo Fisher Scientific, P1304MP) (500 nM, final concentration) and NucBlue Live ReadyProbes Reagent (Hoechst 33342) (Thermo Fisher Scientific, R37605) (5 μg/ml, final concentration) prepared in culture medium were added to the cell media for live staining. The cells were incubated for 30 min with propidium iodide/Hoechst 33342, then the media changed to 10% FBS in HEPES/MEM without phenol red for imaging on a Zeiss confocal microscope (Model LSM 780; Carl Zeiss Microscopy).

#### ER stress assay

The Green/Red Ratiometric Cell Stress Assay (Montana Molecular, U0901G) was used to measure ER stress through fluorescence detection of XBP1 splicing following manufacturers’ protocol. Cells were treated with vehicle (DMSO) or 4 μM 1-deoxysphinganine for 48 h, then imaged under the LSM 780 Zeiss confocal microscope. The images were analyzed for expression of red and green fluorescence intensity using Fiji/ImageJ (37). More than 30 images from each group were analyzed for statistical comparison.

#### Statistical analysis

GraphPad Prism (v.9, GraphPad Software, San Diego, CA) was used for graphing and statistical analyses with one-way ANOVA and unpaired t-tests. All data are presented as mean ± SD. P < 0.05 was considered to be statistically significant and the P values are indicated by asterisks in the figures.

## Data availability

Data generated or analyzed during this study are included in this article. Any additional data is available from the corresponding author upon reasonable request.

## Supporting information

This article contains supporting information.

## Acknowledgments

*—* Supported in part by the Lipidomics Shared Resource, Hollings Cancer Center, Medical University of South Carolina (P30 CA138313 and P30 GM103339). Some Figures created using BioRender.com.

## Funding and additional information

*—* This research was supported by the Intramural Research Program of the National Institute of Diabetes and Digestive and Kidney Diseases (NIDDK) within the National Institutes of Health (NIH). The contributions of the NIH authors are considered Works of the United States Government. The findings and conclusions presented in this paper are those of the authors and do not necessarily reflect the views of the NIH or the U.S. Department of Health and Human Services.

## Abbreviations

—The abbreviations used are:

EMEM: Eagle’s Minimum Essential media;
ER: endoplasmic reticulum;
FBS: fetal bovine serum;
GeCKO: genome-scale CRISPR/Cas9 knockout;
HSAN1: Hereditary Sensory and Autonomic Neuropathy Type 1;
KO: knockout;
LOF: loss-of-function;
MAGeCK: Model-based Analysis of Genome-wide CRISPR/Cas9 Knockout;
RRA: robust ranking aggregation;
S1P: sphingosine-1-phosphate;
SD: standard deviation;
sgRNA: single guide RNA;
SPT: serine palmitoyltransferase;
UPR: unfolded protein response;
XBP1: X-box binding protein 1;
XBP1s: spliced form of X-box binding protein 1.

