## Supplementary material for "1-Deoxysphingolipids Require Very-Long-Chain Ceramide Synthesis to Induce ER Stress and Neurotoxicity": Table S1

**Supplemental Information:**

**Table S1**. **Top 30 positively ranked genes derived from 1-deoxysphinganine resistance screen in SH-SY5Y cells analyzed by Model-based Analysis of Genome-wide CRISPR/Cas9 KO (MAGeCK).**

| id | num | pos\|score | pos\|rank | pos\|lfc |
|---|---|---|---|---|
| *TECR* | 6 | 9.49E-18 | 1 | 6.7474 |
| *ACACA* | 6 | 1.28E-11 | 2 | 3.4526 |
| *CERS2* | 5 | 7.78E-11 | 3 | 3.0367 |
| *PTPLB* | 4 | 2.18E-09 | 4 | 3.7038 |
| *HSD17B12* | 5 | 2.70E-07 | 5 | 0.90363 |
| *hsa-mir-5698* | 1 | 3.38E-05 | 6 | 12.358 |
| *ACLY* | 2 | 0.00060904 | 7 | 2.1624 |
| *HLCS* | 3 | 0.00069476 | 8 | 0.90363 |
| *hsa-mir-3609* | 3 | 0.00069476 | 9 | 0.90363 |
| *ELOVL1* | 3 | 0.00069476 | 10 | 0.90363 |
| *NDUFS8* | 1 | 0.00071066 | 11 | 5.8485 |
| *SRGAP1* | 1 | 0.0010491 | 12 | 5.0126 |
| *SLC46A2* | 1 | 0.0011168 | 13 | 4.9548 |
| *hsa-mir-4278* | 1 | 0.0011844 | 14 | 4.8318 |
| *FAM3D* | 4 | 0.0011966 | 15 | -0.72803 |
| *hsa-mir-3677* | 1 | 0.0013875 | 16 | 4.6916 |
| *PSPH* | 1 | 0.0015567 | 17 | 4.2935 |
| *DHRS7C* | 2 | 0.0016237 | 18 | 0.42207 |
| *WBP5* | 1 | 0.0017259 | 19 | 4.1971 |
| *CCDC101* | 1 | 0.0018274 | 20 | 4.0938 |
| *KAAG1* | 1 | 0.0020981 | 21 | 3.7305 |
| *LHX8* | 1 | 0.0020981 | 22 | 3.7305 |
| *SLC2A11* | 1 | 0.0020981 | 23 | 3.7305 |
| *PLA2G3* | 2 | 0.0026379 | 24 | 1.9105 |
| *RNF144A* | 1 | 0.0026734 | 25 | 3.4249 |
| *FGF16* | 3 | 0.0029413 | 26 | -2.9191 |
| *DYNLL2* | 1 | 0.0031472 | 27 | 3.2438 |
| *KLF1* | 1 | 0.0031472 | 28 | 3.2438 |
| *TACSTD2* | 1 | 0.0031472 | 29 | 3.2438 |
| *SSUH2* | 1 | 0.0031472 | 30 | 3.2438 |
